# Automated monitoring of animal behaviour with barcodes and convolutional neural networks

**DOI:** 10.1101/2020.11.27.401760

**Authors:** Tim Gernat, Tobias Jagla, Beryl M. Jones, Martin Middendorf, Gene E. Robinson

## Abstract

Barcode-based tracking of individuals revolutionizes the study of animal behaviour, but further progress hinges on whether specific behaviours can be monitored. We achieve this goal by combining information obtained from the barcodes with image analysis through convolutional neural networks. Applying this novel approach to a challenging test case, the honeybee hive, we reveal that food exchange among bees generates two distinct social networks with qualitatively different transmission capabilities.

## Main text

Barcode-based tracking makes it possible to automatically identify hundreds of individuals in digital videos, and to record their location and heading direction over long time periods at a high spatiotemporal resolution^1–4^. It generates a wealth of individualized data that transforms how ethologists study the behaviour of animals, especially for species that naturally interact in large collectives, such as ants and honeybees^4–8^. However, in addition to knowing where individuals are located, it is often necessary to also know what they are doing, in order to understand individual or group-level behaviour and its neural and molecular underpinnings.

With the exception of locomotion, barcodes are unable to automatically generate behavioural information directly. When studying the behaviour of barcoded individuals, researchers therefore resort to proxies that infer coarse-grained behavioural states from changes in the location and orientation of an individual’s barcode, and social interactions from the relative position of individuals to each other^2–7,9^ (but see refs. ^1,8^ for two notable exceptions). Such proxies have a limited capacity for distinguishing specific behaviours^10^ and may thus result in high error rates for a particular behaviour of interest.

Convolutional neural networks (CNNs) are a promising technology for developing detectors for specific behaviours. They can be trained to accurately identify digital images that show a particular object, and learn independently which features of the object are most diagnostic^11^. To date, CNNs have been used for animal pose estimation^12–14^ and for detecting behaviours performed in isolation or in small groups. However, localizing behaviour occurrences in an image showing hundreds of individuals continues to be a challenge, especially if the image regions showing the behaviour of interest are small, animals are densely crowded together, or individuals partially occlude each other^15^.

We present a method that combines CNNs with barcode-based tracking to accurately identify specific behaviours in large animal collectives. Our key innovation lies in combining information obtained from an animal’s barcode, such as its location and orientation, with domain knowledge about the behaviour of interest to perform a precise yet computationally inexpensive region proposal that acts as an attention mechanism for the CNN. In addition, we leverage barcode information to simplify the behaviour classification task by rotating the proposed image regions to correspond to a predefined reference frame. Using this approach, we designed barcode-and-CNN-based detectors for two honeybee behaviours, trophallaxis and egg-laying, to demonstrate the power of the method and to show that it can be applied to very different behaviours.

Honeybee trophallaxis is an important social behaviour during which two bees touch each other with their antennae while orally transferring liquid food^16^ and signalling molecules^17^. This behaviour is challenging to detect automatically because honeybee colonies consist of tens of thousands of individuals, and even small experimental colonies still contain hundreds of individuals densely crowded together (Fig. 1a). Moreover, owing to the small size of the honeybee, the detector needs to focus on millimetre-sized body parts, like the mouthparts, to distinguish trophallaxis from visually similar behaviours, such as antennation.

**Figure 1.**
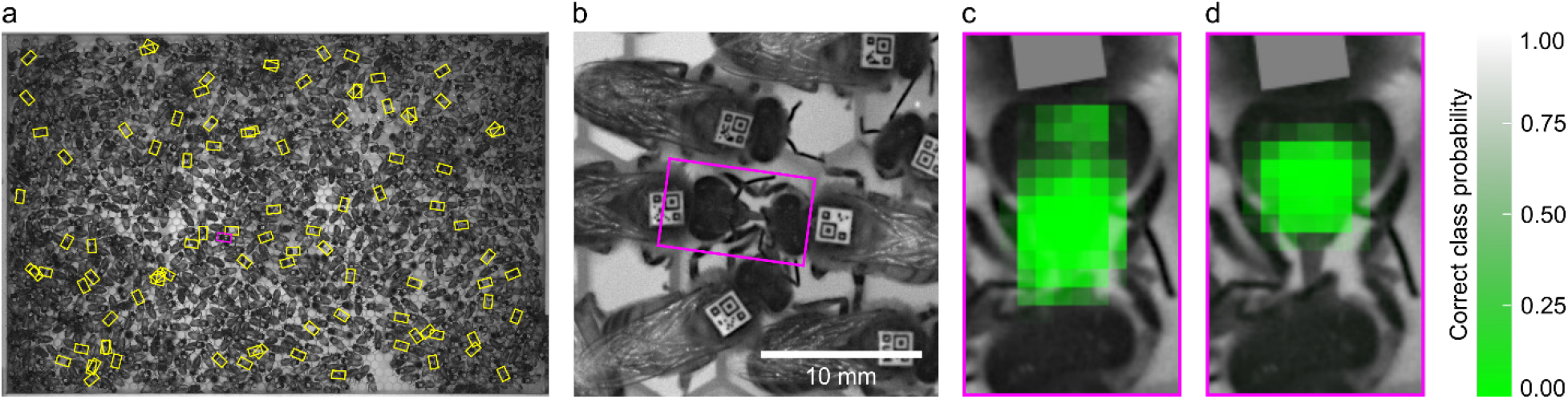
Automatic trophallaxis detection and recipient identification. **a,** Typical image captured by the honeybee tracking rig, showing barcoded bees inside the observation hive. Rectangles outline image regions identified by our barcode-based region proposal procedure for trophallaxis. **b,** Zoom-in on the magenta region proposal in **a**, which shows the head, forelegs and part of the thorax of two bees that engage in trophallaxis. **c**, CNN input image created from the region proposal in **b**, superimposed with a map of the output of the CNN for detecting the occurrence of trophallaxis, as a function of the position of a grey square that occludes some pixels (see Methods). The input image is more likely to be misclassified (low probability of trophallaxis) if pixels corresponding to the mouthparts and proboscis are obscured. This result indicates that the CNN is able to distinguish salient features of trophallaxis from the background. **d**, The same input image as in **c**, but superimposed with a map of the correct class probability for recipient identification. This map suggests that the CNN for identifying the recipient relies on visual cues obtained from the mouthparts of the recipient, which, unlike those of the donor, are always visible and closely aligned with the proboscis.

Our trophallaxis detector partitions recognizing the occurrence of this behaviour into four tasks: region proposal, preprocessing, visual verification, and post-processing (see Methods). Region proposal first selects pairs of bees that are in the proper position relative to each other to orally exchange liquid food (Supplementary Fig. 1). It then precisely estimates the image region that shows the mouthparts, proboscis (“tongue”), and head of the potential trophallaxis partners (Fig. 1a,b). These two steps exclude 97.9±1.0% (mean ± standard deviation, n=300) of an image from automatic visual examination, which increases the detector’s computational efficiency and reduces its false positive rate. Each proposed region is then preprocessed and scored with a CNN that has been trained to estimate the probability that the region shows bees engaging in trophallaxis (Supplementary Table 1 and Fig. 1c).

Performance measurements showed that our barcode-and-CNN-based trophallaxis detector significantly outperforms the state of the art (Supplementary Table 2). Its Matthews correlation coefficient (MCC) is 0.89, and thus 0.28 higher than that of the best automatic trophallaxis detector so far^8^. Most of this improvement is due to a 67% higher sensitivity and an 11% higher positive predictive value, which means that our detector identifies more trophallaxis interactions and generates fewer false positives. Unlike existing automatic trophallaxis detectors^8,9^, it is also able to accurately score interactions between workers and the queen, even though social interactions with the queen differ from worker-worker interactions in that she is constantly antennated and groomed^16^. Moreover, at 57.7 megapixel/s our trophallaxis detector processes digital images 16.7 times faster than the reference detector^8^, enabling larger-scale experiments.

When studying social behaviour, it is often important to know the role played by each individual, i.e., who is the “donor” and who is the “recipient” of a particular behavioural interaction. To obtain this information for trophallaxis, we trained a second CNN (Supplementary Table 1 and Fig. 1d) to identify the liquid recipient, and operated it in parallel with the CNN that detects the occurrence of trophallaxis. Previously described automatic trophallaxis detectors did not attempt to determine the direction of liquid flow^8,9^, which involves distinguishing the individual that has only opened her mouthparts (donor) from the one that has also extended her proboscis (recipient). When applied to automatically identified trophallaxis partners, our recipient detector has a MCC of 0.97.

To verify that our trophallaxis detector generates plausible results, we used it to monitor trophallaxis in three honeybee colonies, consisting of up to 1,050 barcoded individuals, at 1 frame/s for five consecutive days. We then employed an epidemiological model to perform bidirectional spreading simulations on the trophallaxis networks of these colonies (Supplementary Table 3). As these simulations ignore the direction of liquid flow, they can be used to study information and disease transmission via physical contacts that take place during trophallaxis, but not how liquid flows through the trophallaxis network. Consistent with previous results^8^, we observed that bidirectional spreading through the observed trophallaxis networks was faster than through their temporally randomized counterparts (Fig. 2, Supplementary Figs. 2 and 3), confirming that physical contacts during trophallaxis have the ability to accelerate the transmission of information or disease.

**Figure 2.**
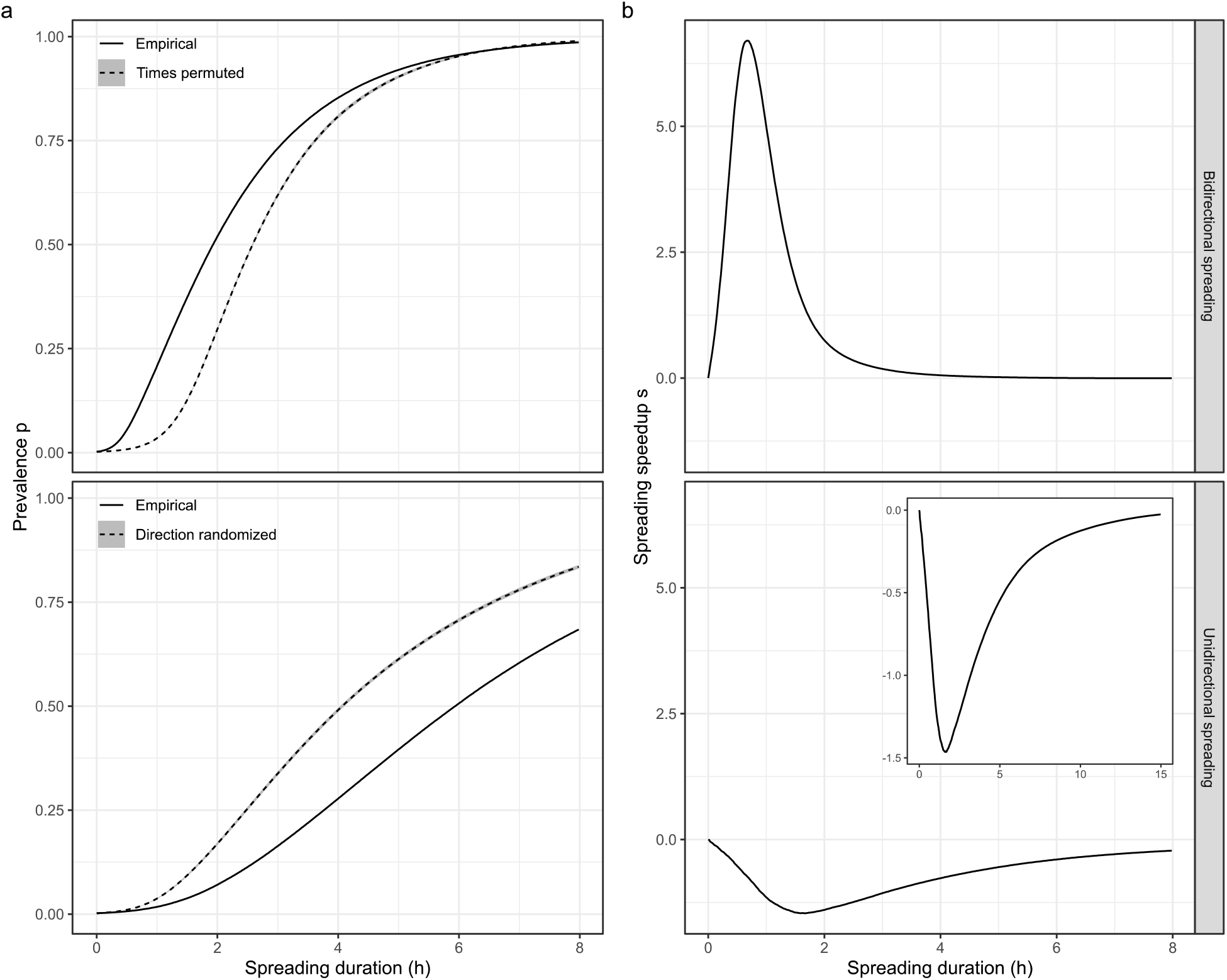
Simulated spreading though honeybee trophallaxis networks. Panels show data from Trial 1; see Supplementary Figs. 2 and 3 for Trial 2 and 3, respectively, which yielded similar results. Top row: bidirectional spreading results, modelling to transmission via physical contacts during trophallaxis; bottom row: unidirectional spreading results, modelling liquid transfer. **a,** Average fraction of “infected” bees (prevalence) as a function of spreading duration. Solid line: prevalence in the empirical trophallaxis network; dashed line: mean prevalence, averaged across 10 randomized reference networks; grey band: point-wise 95% confidence interval. Simulation results indicate that the temporal pattern of trophallaxis accelerates bidirectional spreading, and that the directional nature of trophallaxis inhibits unidirectional spreading. **b,** Spreading speedup, corresponding to the normalized difference between the prevalence in the empirical network and the mean prevalence across the 10 randomized reference networks, as a function of spreading duration. The spreading speedup is positive if the prevalence in the empirical trophallaxis network is higher than in the randomized reference networks, zero if there is no difference, and negative if it is lower. Inset: Spreading speedup as a function of spreading duration until, like for bidirectional spreading, almost all bees are “infected”, showing that the spreading speedup remains negative.

Automatic identification of the recipient allowed us for the first time to also simulate unidirectional spreading dynamics. These simulations take into account that liquid gets transferred from the donor to the recipient (see Methods), and thus model the flow of liquid through the trophallaxis network. By comparing unidirectional spreading dynamics between the observed trophallaxis networks and directionally randomized reference networks, we discovered that the simulated liquid flow is slower than expected by chance (Fig. 2, Supplementary Figs. 2 and 3). This result suggests that the directional nature of trophallaxis serves to inhibit the transmission of liquids and the compounds they contain. It furthermore suggests that even though the network of physical contacts during trophallaxis and the network of actual liquid transfers result from the same behaviour, they are distinct entities with qualitatively different transmission properties, which may have important implications for the function and organisation of honeybee colonies.

To show that our approach to automatic behaviour detection generalizes to other behaviours, we developed a barcode-and-CNN-based detector for egg-laying (see Methods), a solitary behaviour performed by worker honeybees when the colony has lost its primary reproductive, the queen, and is unable to replace her. Because we included no proxy for identifying bees that are likely to lay an egg in the region proposal procedure for this detector, it is applied to all bees in an image and therefore generated more false positives. We addressed this problem by adding to this detector a second CNN that was trained to identify and weed out false positives generated by the first CNN. This approach increased the detector’s MCC from 0.64 to 0.76 (see Supplementary Table 2 for additional performance values), demonstrating that barcode-and-CNN based detectors can identify visually very different behaviours.

Automatically detecting animal behaviour is difficult. We showed that barcode-and-CNN-based detectors can accomplish this task even under challenging conditions, if domain knowledge and information obtained from an individual’s barcode are used to simplify the behaviour classification task. Our approach can be applied to all animals that perform visually recognizable behaviours, and to which a trackable marker can be affixed. We therefore envision that barcode-and-CNN-based detectors will make it possible to monitor behaviour in a variety of species, and thus pave the road for automatic, high-resolution behavioural studies that address a broad range of previously intractable questions in ethology, neuroscience, and molecular biology through quantitative analysis of specific behaviours.

## Methods

### Honeybee tracking

Colonies were established as in ref. ^8^. Briefly, up to 1,370 one day old honeybee workers were individually outfitted with a barcode by chilling them until they stopped moving and then gluing a bCode barcode to their thorax. For recording trophallaxis interactions, we also barcoded an unrelated, naturally mated queen.

All barcoded bees passing quality control were moved into an observation hive, which was placed in a dark, climate-controlled room and connected to the outside environment via a plastic tube to enable normal foraging. The observation hive held a 348 mm × 232 mm white plastic honeycomb, one side of which was inaccessible to the bees. The other side was provisioned with enough honey and pollen for the duration of the experiment, and covered by an exchangeable glass window. The distance between this window and the honeycomb was short enough to ensure that the bees formed a monolayer, which does not affect their behaviour^8^ and prevents them from obscuring their barcodes.

Trophallaxis interactions and egg-laying events were monitored with the tracking systems described in ref. ^18^ (Supplementary Fig. 4) and ref. ^8^, respectively. Video of the honeycomb was captured for up to 7 consecutive days at a frequency of 1 frame/s with a computer-controlled 29 megapixel monochrome machine vision camera under infrared light, invisible to the bees, and stored on hard drives for later processing. Barcode detection and decoding was performed on a compute cluster, using the software and procedures described in ref. ^8^.

### Ground truth

To create a ground truth for the trophallaxis detector, we manually annotated images of pairs of bees that were in the proper position to perform this behaviour. For each image we recorded whether the two bees were engaged in trophallaxis and, if so, which bee was the recipient.

The first set of images consisted of the image library *L1* described in ref. ^8^. These images show random bee pairs that were selected solely based on whether they were within reach. To determine if two bees are within reach, the coordinate of their mouthparts was estimated by translating the barcode centre of each bee by a fixed distance in the direction of her barcode orientation vector, which was assumed to be parallel to her anteroposterior axis (Supplementary Fig. 1). If the distance between these coordinates was shorter than 7 mm, which is the maximum proboscis length of a honeybee^19^, the image was annotated. An image was considered to show trophallaxis if the proboscis of one bee touched the head of the other bee close to her mouthparts (Fig. 1b). Otherwise it was annotated as not showing trophallaxis.

Examination of the images in library *L1* revealed that only approximately 1 in 40 images showed trophallaxis. Because CNNs learn to classify images better if the number of positive and negative examples is more balanced^20^, and manual annotation is labour intensive, additional images were selected by also requiring that bees needed to face each other. Whether two bees faced each other was established similar to ref. ^8^: We calculated the angle between the barcode orientation vector of each bee and a line through the mouthpart location of both bees (Supplementary Fig. 1). If the sum of the two resulting angles was less than 104 degrees, the image was annotated. This criterion captures 95% of the trophallaxis contacts in *L1*. Using this filter as a proxy for identifying trophallaxis, we annotated an additional 6045 random bee pair images.

To evaluate the performance of the trophallaxis detector when its predictions are integrated over time, we furthermore used the trophallaxis proxy to annotate all bee pairs in 600 random triples of successively recorded images of the entire observation hive. Each bee pair was annotated by 3 annotators to be able to reduce annotation errors. Consensus among the 3 annotators was reached through majority voting. A subset of the resulting annotations were previously published as image library *L2*^8^.

The ground truth for the egg-laying detector was created by manually annotating 1323 random images of the entire hive. Bees that either had inserted their abdomen into a honeycomb cell (Supplementary Fig. 5) or appeared to be in the process of doing so were called egg-layers. All other bees were annotated as having laid no egg. In addition we annotated each egg layer in up to two images that were recorded before and after the “focal” hive image.

The final trophallaxis and egg-laying ground truths consisted of 142,182 images of bee pairs and 723,995 images of individual bees, respectively. Both ground truths were split into disjunct training, calibration, and test data sets as shown in Supplementary Table 6.

### Region proposal

To identify image regions that are likely to show bees engaged in trophallaxis, we leveraged the trophallaxis proxy described earlier (Supplementary Fig. 1). For each pair of potential trophallaxis partners identified by this proxy, we extracted the region showing the heads of both bees by translating the barcode centre of each bee by a fixed distance in the direction of her barcode orientation vector. The midpoint of the line segment defined by these two coordinates was used as the centre point of a 96 px × 160 px region of which the longer sides were parallel to the aforementioned line segment (Fig. 1b). The top edge of this rectangle was defined to be the short edge closest to the head of the bee with the bigger ID.

For detecting egg-layers, the image region focusing on the abdomen of a bee was extracted by translating the centre of a bee’s barcode by a fixed distance in the opposite direction of her barcode orientation vector. The resulting coordinate was used as the midpoint of one edge of a 130 px × 130 px region of which the top edge was parallel and closest to the lower edge of the bee’s barcode (Supplementary Fig. 5a). The image region showing the entire bee as well as her immediate surroundings was obtained by first translating the centre of a bee’s barcode by a fixed distance in the direction of her barcode orientation vector. The resulting coordinate was then used as the top-right corner of a 256 px × 256 px region of which the diagonal between the top-right corner and the bottom-left corner passed through the barcode centre (Supplementary Fig. 5b).

### Image preprocessing

CNN input images were created by extracting the proposed image region and rotating it “upright”, so its top edge was on the x-axis of the image coordinate system. Pixel intensities were then clamped at a value of 200 to remove bright details in the background that we did not expect to provide information about the behaviour of interest, such as the honeycomb structure and specular reflections in the honeycomb cell contents (Fig. 1c). For trophallaxis detection, we furthermore filled the bounding box of the focal bees’ barcode with a uniform colour to prevent the CNN from associating parts of the barcode pattern with a behaviour (Fig. 1c). Finally, pixel intensities where mean-centred and scaled to [-1, 1] to provide a consistent input range for the CNNs.

### CNN architecture and training

Images of potential trophallaxis partners and egg-layers were classified with two CNNs, each. While the exact details of these CNNs varied, they had a similar architecture, consisting of 2 or 3 convolutional layers, max-pooling layers, and 2 fully connected layers (Supplementary Tables 1, 4, and 5). The output of convolutional layers was standardized with batch normalization before being passed to a rectified linear unit activation function, and the output of the final activation function was transformed by a softmax function, so it can be interpreted as the probability of the input image showing the behaviour of interest.

CNN training consisted of initializing the weights of the network to values drawn from a normal distribution with a mean of 0 and a standard deviation of 1, truncated at 2 standard deviations. Network weights were then optimized for 10,000 iterations, using the Adam algorithm, which was configured as recommended by its authors (alpha=0.001, beta1=0.9, beta2=0.999, epsilon=0.0000001)^21^, with a cross-entropy loss function. We used batches of 256 images, which were augmented as shown in Supplementary Table 7 and normalized to mean zero and unit variance. To avoid overfitting, we applied a L2 weight decay of 0.005 and a dropout of 0.5 in the first fully connected layer.

The CNNs for detecting the occurrence of trophallaxis and whether a bees’ abdomen was inserted into a honeycomb cell were trained on all positive examples and a matching number of random negative examples from the trophallaxis and egg-laying training data set, respectively. For training the CNN that identifies the recipient, we used all positive examples of the trophallaxis training data set and a corresponding number of random negative examples. The CNN that uses an image of the entire bee to identify false positives was trained on the 4044 false positives generated by the CNN that checks for the visual absence of a bee’s abdomen and a matching number of random positive examples from the egg-laying training data set.

### Behaviour detection

To detect trophallaxis or egg-laying in individual images, we first performed the corresponding region proposal to obtain image regions that potentially show the behaviour of interest. Each of these regions was extracted, preprocessed as described above, and independently scored by the two CNNs that had been trained to detect the behaviour. This approach is computationally less efficient than a strictly serial operation where, for example, the CNN for identifying the recipient processes the input image only if the score for the occurrence of trophallaxis exceeds a specified threshold. It was nevertheless chosen because it makes it possible to store both CNN scores in a file, which can later be post-processed to yield behaviour predictions of varying stringency without having to process the video again.

Successive per-image trophallaxis and egg-laying detections were thresholded and linked together to yield behaviour predictions that are easier to analyse. Linked predictions were filtered to improve their quality. For trophallaxis, this procedure is identical to the post-processing steps described in ref. ^8^. Briefly, detections between the same two individuals were concatenated if they occurred in successive video frames. Concatenated detections shorter than 3 s were discarded, because such short trophallaxis interactions may not always result in the transfer of liquid^22^. The remaining interactions were merged if they were less than 60 s apart, and discarded if their total duration exceeded 180 s.

For egg-laying, linking and filtering consisted of concatenating the thresholded detections from successive video frames into egg-laying events. Concatenated egg-laying events shorter than 3 s were discarded. The remaining events were merged if they involved the same bee, occurred within 10 s of each other, and the distance of the average position of the events was shorter than 11.2 mm (the width of two honeycomb cells). These conditions ensured that egg-laying predictions were only merged if they appeared to belong to the same real event.

### Detector calibration

The trophallaxis detector and egg-laying detectors each have free parameters. For the trophallaxis detector, these parameters are the minimum and maximum distance between a pair of bees, the maximum sum of the angle between the bees’ orientation vector and a line through their estimated mouthparts location, a threshold for the output of the CNN that determines which scores correspond to the occurrence of trophallaxis, and a threshold for the output of the CNN that identifies the recipient. For the egg-laying detector, free parameters are the CNN thresholds that determine which output scores identify a (potential) egg layer.

To fix the free parameters of each behaviour detector, we applied it to the calibration data set of its ground truth. We then performed a grid search on the parameter space of the detector, and chose the parameter combination that maximized the product of the detector’s sensitivity and positive predictive value.

### Evaluation

We estimated the fraction of an image that the trophallaxis detector’s region proposal procedure excludes from automatic visual inspection by applying it to the “centre” image of all 300 observation hive image triples from which the detector’s test data set was created. We then added the number of pixels inside the proposed image regions and divided the total by the number of pixels per image and by the number of observation hive images.

To test if the CNNs for detecting the occurrence of trophallaxis and for identifying the recipient are sensitive to salient features of the behaviour, we performed an occlusion sensitivity analysis^23^. More specifically, we systematically moved a 41 px × 41 px big grey square (occluder) over the CNN input image, restricting the occluder centre to pixels inside the image. For each occluder position, we processed the altered input image with the respective CNN and recorded the CNN’s output. CNN outputs were then spatially coarse-grained into 12 × 20 square bins by averaging over all outputs inside a bin. Bins with a low mean score represent occluder positions that lead to a misclassification of the altered input image.

Detector performance was evaluated by applying each detector to its respective test data set, using the parameters we had obtained during detector calibration. This results in an estimate of a detector’s performance on images of the entire hive, since the prevalence of trophallaxis and egg-laying in the respective test data sets is the same as in hive images. These performance estimates are conservative, because the test data sets consist of 3 s long segments of behaviour occurrences that likely lasted longer. Longer behaviour occurrences consist of multiple such segments and are therefore more likely to be detected. Moreover, due to the high positive predictive value of both detectors on individual images, the probability of a spurious detection decreases sharply as the duration of the detected behaviour increases.

Image processing time was measured by averaging detector runtime across the 900 (545) hive images from which the trophallaxis (egg-laying) test data set was created. These measurements were performed on a cluster with 2.9 GHz Intel Core i9-7920X CPUs and a RAID 6 storage array consisting of 6 HGST H3IKNAS800012872SWW hard drives. During runtime measurements, both detectors were restricted to a single hardware thread and had access to 2 GB RAM.

### Spreading simulations

To simulate the transmission of information, pathogens, and liquid, we employed a temporally explicit version of the deterministic susceptible-infected model^24^. This model assumes that individuals are in one of two states, “susceptible” or “infected”. Simulations begin by setting all bees to susceptible, choosing a trophallaxis interaction uniformly at random, and “infecting” the two bees involved in this interaction. Spreading dynamics were then simulated over an 8 h time window. During this time, an infected donor “infects” a susceptible recipient with a probability of 1 when they engage in trophallaxis (unidirectional transmission). For models of bidirectional transmission, the trophallactic role of individuals was ignored, which means recipients were also able to infect donors.

For each simulation, we recorded the fraction of infected individuals*f*(*f*)=*i*(*t*)/*S*(*t*), where *i*(*t*) is the number of infected bees alive at time *t* after the first infection, and *S*(*t*) is the colony size at time *t*, both of which are subject to mortality. To obtain a more robust estimate of fraction of infected individuals, we averaged the fraction of infected individuals over *R*=1,000 simulation runs and calculated the prevalence 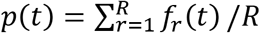.

To establish whether the prevalence for an observed interaction sequence, 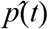, is greater than expected by chance, we compared it to the prevalence for *N*=10 randomized interaction sequences. For bidirectional spreading simulations, randomized interaction sequences were created with the PTN null model^25^, which shuffles the contact times among the observed interactions without artificially prolonging an individual’s life. For unidirectional spreading simulations, interaction sequences were randomized by reversing the direction of trophallaxis with a probability of 0.5. Spreading dynamics where characterized by calculating the spreading speedup 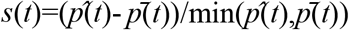, where 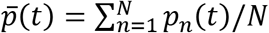 is the mean prevalence across the *N* randomized interaction sequences.

## Acknowledgements

We thank the University of Illinois School of Life Sciences Machine Shop for constructing bee tracking equipment; Reliance Label Solutions for printing bCodes; and the Carl R. Woese Institute for Genomic Biology Computer Network Resource Group for computational support. We are grateful to J. Peng for providing computational resources. We thank J. Cullum, S. Bransley, and A. Ray for manual image annotation; T. L. Harrison and A. L. Sankey for bee management; and A. R. Hamilton, A. Ray, S. Bransley, J. Cullum, K. Wilk, J. Falk, A. Zhang, L. Block, and V. Bagchi for assistance with field work. We furthermore thank members of the M.M. and G.E.R. laboratories for discussions; and members of the M.M. and G.E.R. laboratory, Jan Aerts, and our anonymous reviewers for comments that improved the manuscript. This material is based on work supported by the National Academies Keck Futures Initiative Grant NAKFI CB4 (to T.G.), a grant from the Christopher Family Foundation (to G.E.R.), National Institutes of Health Grant R01GM117467 (to G.E.R. and Nigel Goldenfeld), Defense Advanced Research Projects Agency Gant HR0011-16-2-0019 (to G.E.R. and Huimin Zhao), and National Institute of General Medical Sciences Grant R01GM117467 (to G.E.R. and Nigel Goldenfeld).

---

**Supplementary Figure 1.**
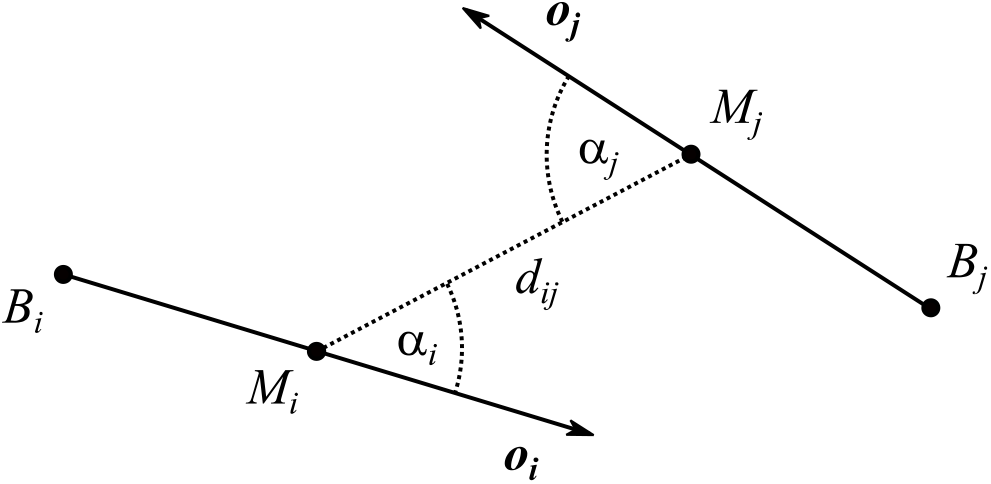
Spatial proxy for identifying pairs of bees that might be engaged in trophallaxis. Points *B_i_* and *B_j_* are the barcode centres of bee *i* and *j*, respectively. Arrows represent the barcode orientation vectors ***o**_i_* and ***o**_j_* that correspond to the heading direction of a bee. Points *Mi* and *M_j_* are the estimated mouthparts locations of the two bees, and *d_ij_* is the distance between these points. If *d_ij_* was shorter than the proboscis length of a honeybee and the sum of the angles *α_i_* and *α_j_* is smaller than a specified threshold, bees *i* and *j* were called potential trophallaxis partners.

**Supplementary Figure 2.**
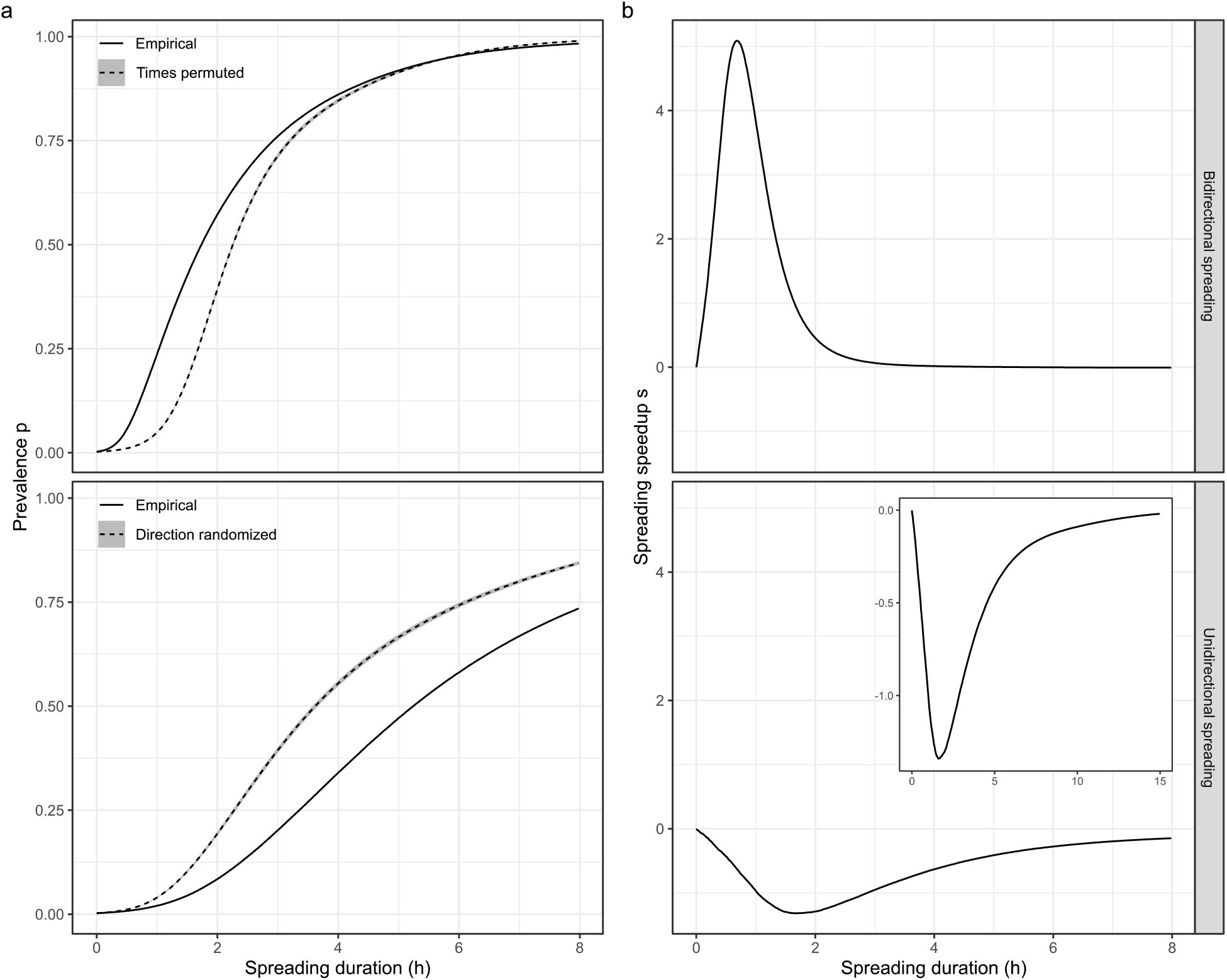
Simulated spreading in honeybee trophallaxis networks. Panels show data from Trial 2. Top row: bidirectional spreading results, modelling to transmission via physical contacts during trophallaxis; bottom row: unidirectional spreading results, modelling liquid transfer. **a,** Prevalence as a function of spreading duration. Solid line: prevalence in the empirical trophallaxis network; dashed line: mean prevalence, averaged across 10 randomized reference networks; grey band: point-wise 95% confidence interval. **b,** Spreading speedup as a function of spreading duration. Inset: Spreading speedup as a function of spreading duration until, like for bidirectional spreading, almost all bees are “infected”.

**Supplementary Figure 3.**
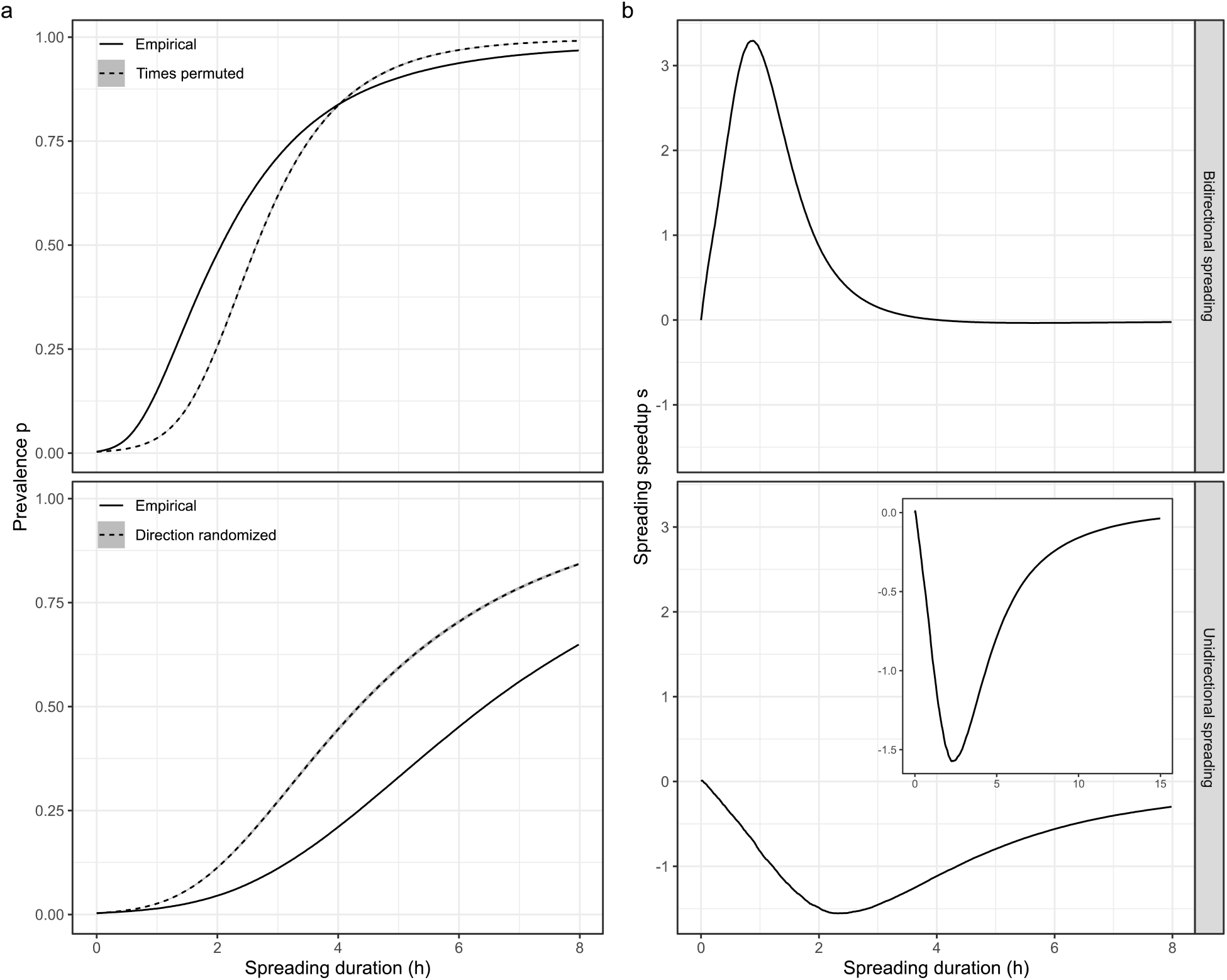
Simulated spreading in honeybee trophallaxis networks. Panels show data from Trial 3. Top row: bidirectional spreading results, modelling to transmission via physical contacts during trophallaxis; bottom row: unidirectional spreading results, modelling liquid transfer. **a,** Prevalence as a function of spreading duration. Solid line: prevalence in the empirical trophallaxis network; dashed line: mean prevalence, averaged across 10 randomized reference networks; grey band: point-wise 95% confidence interval. **b,** Spreading speedup as a function of spreading duration. Inset: Spreading speedup as a function of spreading duration until, like for bidirectional spreading, almost all bees are “infected”.

**Supplementary Figure 4.**
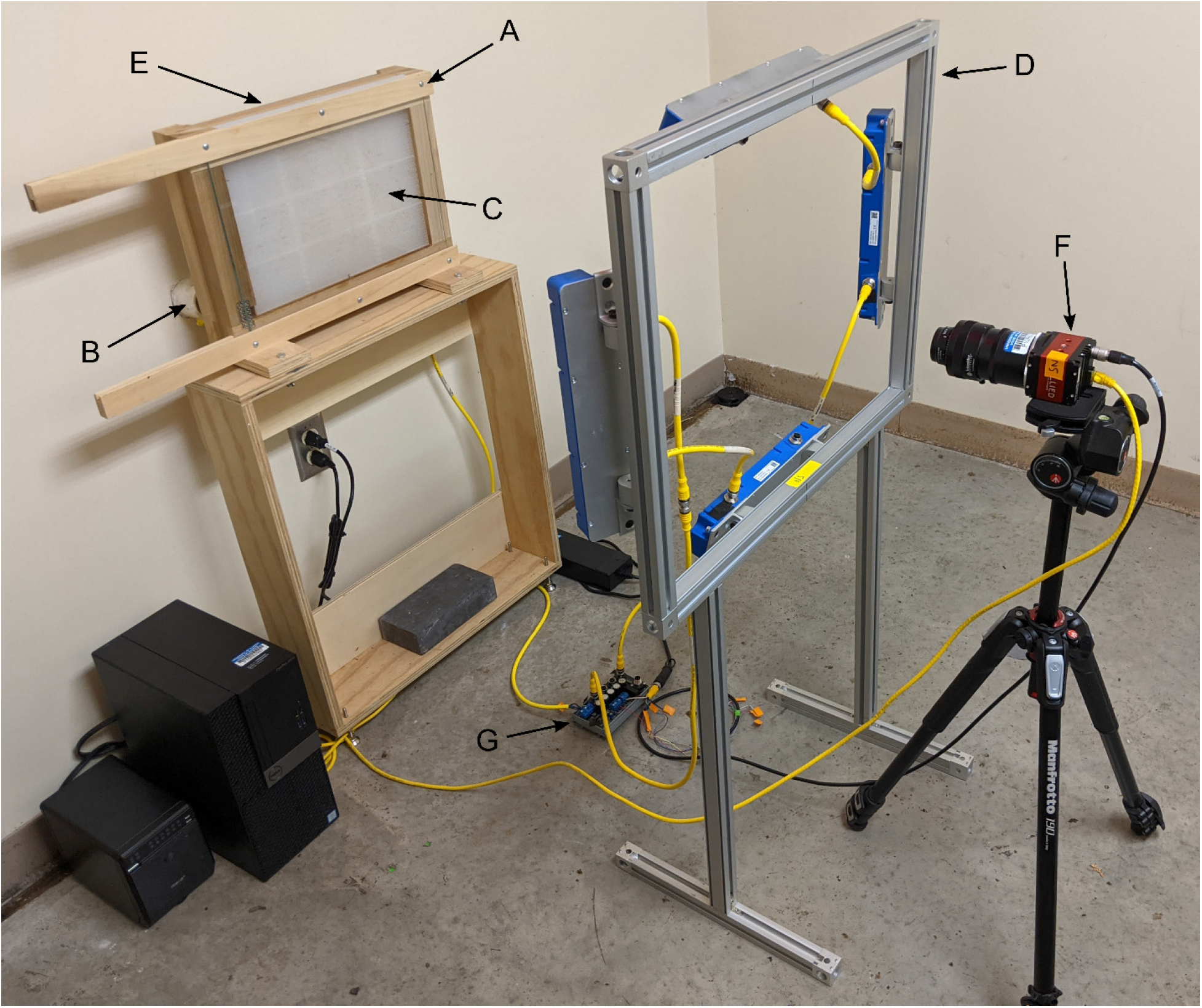
Setup for tracking barcoded honeybees and automatically monitoring their behavior. Barcoded bees were housed in an observation hive (A) that was connected to the outdoors via an entrance tube (B). The hive held a glass-covered, one-sided plastic honeycomb (C), which was front-lit with four infrared LED lights mounted on an aluminum frame (D) and backlit with an array of infrared lights mounted behind the hive (E, hidden). A computer-controlled high-resolution monochrome camera (F) recorded the hive, triggering the infrared lights via a breakout board (G).

**Supplementary Figure 5.**
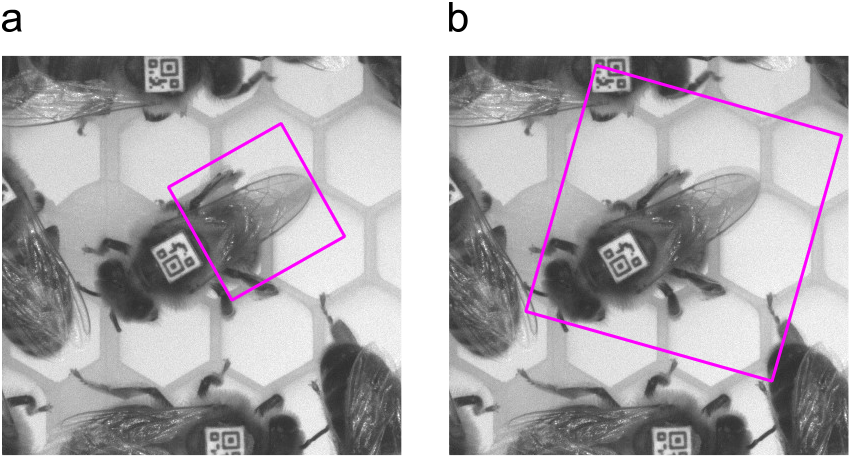
Region proposals for the egg-laying detector. Proposed image regions are shown as magenta rectangles. **a,** Image region focusing on the bee’s abdomen, which is invisible because the bee has inserted it into a honeycomb cell to position an egg. This region was used to identify potential egg-layers. **b,** Image region of an entire bee. This region was used to classify potential egg-layers into true egg-layers and false positives.

**Supplementary Table 1.** Architecture of the CNN for predicting the occurrence of trophallaxis and of the CNN for identifying the recipient. I: input layer; C: convolutional layer; MP: max-pooling layer; F: fully connected layer. Layer output dimensions are W × H × D.

| Layer | Layer type | Kernel size | Stride | Output dimensions |
| --- | --- | --- | --- | --- |
| 0 | I | - | - | $36 \times 60 \times 1$ |
| 1 | C | $5 \times 5$ | 1 | $36 \times 60 \times 8$ |
| 2 | MP | $2 \times 2$ | 2 | $18 \times 30 \times 8$ |
| 3 | C | $3 \times 3$ | 1 | $18 \times 30 \times 16$ |
| 4 | MP | $2 \times 2$ | 2 | $9 \times 15 \times 16$ |
| 5 | F | - | - | $1 \times 1 \times 32$ |
| 6 | F | - | - | $1 \times 1 \times 2$ |

**Supplementary Table 2.** Detailed detection and runtime performance estimates. Note that the runtime estimate for trophallaxis detection includes detecting the occurrence of trophallaxis as well as identifying the recipient.

| **Detector** | **Sensitivity** | **Specificity** | **Positive**<br>**predictive value** | **Negative**<br>**predictive value** | **Speed**<br>**(s/image)** |
| --- | --- | --- | --- | --- | --- |
| Trophallaxis | 0.89 | 1.00 | 0.90 | 1.00 | 0.52 |
| Egg-laying | 0.63 | 1.00 | 0.93 | 1.00 | 1.38 |

**Supplementary Table 3.** Trophallaxis network properties.

| **Trial** | **Nodes** | **Edges** | **Interactions** |
| --- | --- | --- | --- |
| 1 | 1,050 | 115,269 | 150,859 |
| 2 | 902 | 97,591 | 132,591 |
| 3 | 815 | 60,540 | 84,164 |

**Supplementary Table 4.** Architecture of the CNN for predicting whether a bee has inserted her abdomen into a honeycomb cell. I: input layer; C: convolutional layer; MP: max-pooling layer; F: fully connected layer. Layer output dimensions are W × H × D.

| Layer | Layer type | Kernel size | Stride | Output dimensions |
| --- | --- | --- | --- | --- |
| 0 | I | - | - | $32 \times 32 \times 1$ |
| 1 | C | $3 \times 3$ | 1 | $32 \times 32 \times 32$ |
| 2 | MP | $2 \times 2$ | 2 | $16 \times 16 \times 32$ |
| 3 | C | $5 \times 5$ | 1 | $16 \times 16 \times 64$ |
| 4 | MP | $2 \times 2$ | 2 | $8 \times 8 \times 64$ |
| 5 | F | - | - | $1 \times 1 \times 256$ |
| 6 | F | - | - | $1 \times 1 \times 2$ |

**Supplementary Table 5.** Architecture of the CNN for classifying the predictions generated by the CNN shown in Supplementary Table 4 into true egg-layers and false positives. I: input layer; C: convolutional layer; MP: max-pooling layer; F: fully connected layer. Layer output dimensions are W × H × D.

| Layer | Layer type | Kernel size | Stride | Output dimensions |
| --- | --- | --- | --- | --- |
| 0 | I | - | - | $64 \times 64 \times 1$ |
| 1 | C | $5 \times 5$ | 1 | $64 \times 64 \times 32$ |
| 2 | MP | $2 \times 2$ | 2 | $32 \times 32 \times 32$ |
| 3 | C | $3 \times 3$ | 1 | $32 \times 32 \times 64$ |
| 4 | MP | $2 \times 2$ | 2 | $16 \times 16 \times 64$ |
| 5 | C | $3 \times 3$ | 1 | $16 \times 16 \times 128$ |
| 6 | MP | $2 \times 2$ | 2 | $8 \times 8 \times 128$ |
| 7 | F | - | - | $1 \times 1 \times 256$ |
| 8 | F | - | - | $1 \times 1 \times 2$ |

**Supplementary Table 6.**
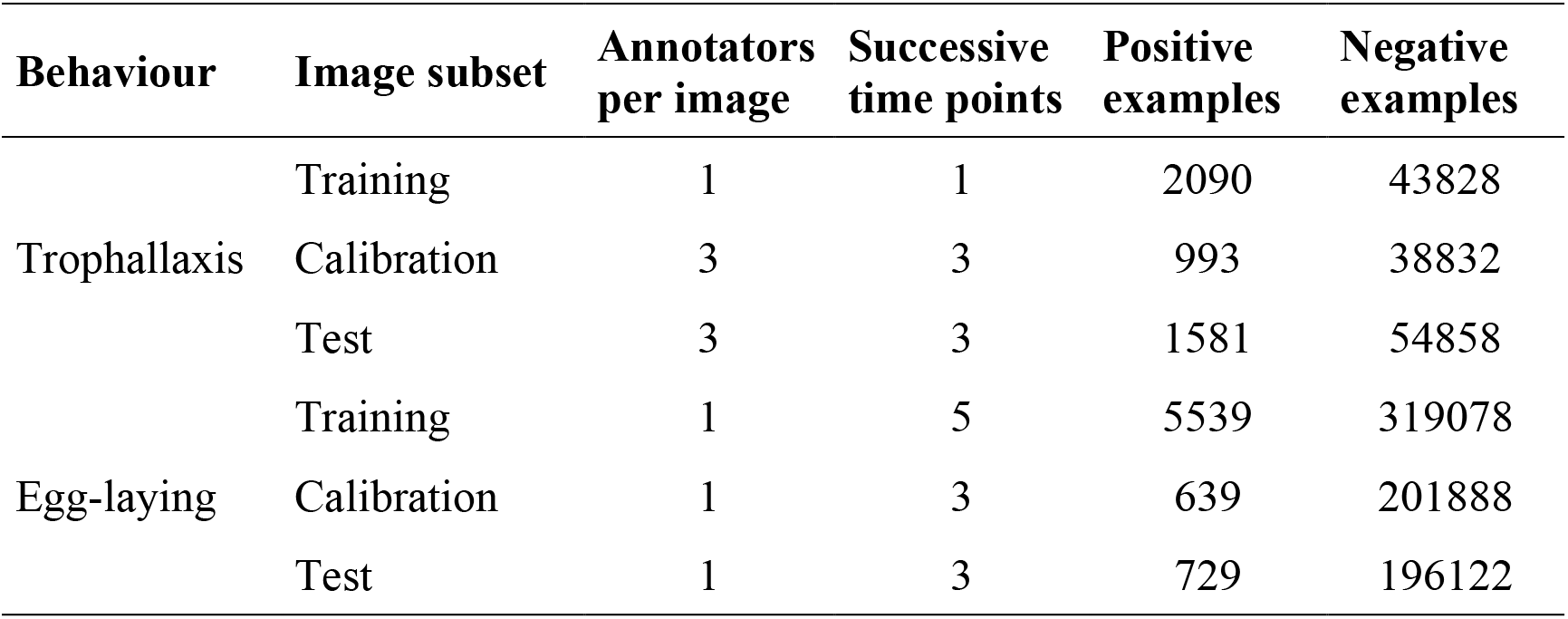
Overview of the trophallaxis and egg-laying gold standards. The number of successive time points indicates if image sequences were annotated, and how long these sequences were. The number of positive and negative examples shows the number of images per subset. The number of annotations per subset is therefore higher than the number of images per subset if multiple annotators scored each image.

**Supplementary Table 7.** Overview of the data augmentation techniques used to increase the diversity of the data sets for training the two CNNs employed by the trophallaxis detector and the egg-laying detector. A technique was applied to each example image with the specified probability if the corresponding table cell shows a plus sign; otherwise the technique was not used.

| Technique | Probability | Trophallaxis |  | Egg-laying |  |
| --- | --- | --- | --- | --- | --- |
|  |  | Trophallaxis detection | Recipient identification | Abdomen detection | Pose examination |
| Adjust brightness | 1 | + | + | + | + |
| Adjust contrast | 1 | + | + | + | + |
| Jitter | 1 | + | + | + | + |
| Flip vertical | 0.5 | + | + | + | - |
| Flip horizontal | 0.5 | + | + | - | - |
| Flip diagonal | 0.5 | - | - | - | + |

